# The Efficiency, Efficacy, and Retention of Motor Training in Chronic Stroke

**DOI:** 10.1101/2020.01.27.922096

**Authors:** Chunji Wang, Carolee Winstein, David Z. D’Argenio, Nicolas Schweighofer

## Abstract

In motor skill learning, the greater the dose of training, the greater the efficacy of training, the lower the efficiency of training, and the better the long-term retention. Whether such principles apply to motor training after stroke is unclear. Here, we developed novel mixed-effects models of the change in the quality of arm movements during and following training fitted to data from a recent randomized controlled trial of the effect of the dose of training in chronic stroke. Analysis of the model’s learning and retention rates demonstrated an increase in efficacy of training with greater doses, a decrease in efficiency of training with both additional doses and additional bouts of training, and fast initial decay following training. Two additional effects modulated retention: a positive “self-training” effect, and an un-expected negative effect of dose. Our results suggest that for patients with sufficient arm use post-training, self-training will further improve use, but additional therapy may be in vain.

## Introductions

Rehabilitation in the chronic phase after stroke is based on the premise that motor learning determines activity-dependent sensorimotor recovery^1–4^. Extensive research in motor skill learning^5^ has shown that: 1) increasing the amount of training increases efficacy (“practice makes perfect”), 2) increasing the amount and duration of training decrease efficiency (“the power law of practice”^6^), and 3) increasing the amount of training improves retention following training. Whether these principles translate to clinical outcomes during and following motor training in stroke survivors, remains unclear for three reasons. First, the effect of increasing the amount of training on clinical outcomes is controversial. While both our recent Dose Optimization for Stroke Evaluation (DOSE) trial and meta-analyses^7, 8^ suggest that higher doses lead to greater gains^9^, a recent phase II clinical trial showed no effect of dose on arm function^10^. Second, although studies of motor training after stroke show that most gains in movement performance are achieved in the initial sessions^11, 12^, the changes of efficiency of training with both additional doses and additional bouts of training on clinical outcomes are unknown. Third, retention following motor training in stroke survivors appears highly variable and the reasons for such variability are unclear. Whereas some studies have shown decay of the gains post-training, as least for sub-groups of patients^13^, others have shown that the gains can be maintained after training^11, 14^, yet others have shown that the gains can even further increase following training, possibly through “self-training” if arm use is sufficient^13, 15–17^

Here, we analyzed the changes in the Quality of Movement sub-scale of the Motor Activity Log (henceforth, the MAL) in the DOSE trial^9^, in which participants with chronic stroke were randomized into four dose groups. The total doses (0, 15, 30 and 60 hours) were equally distributed over the three bouts of training, as shown in Figure 1B. The MAL was collected in 14 longitudinal assessments before and after training in each of the three 1-week training bouts and then monthly for 6 months following the last training bout.

**Figure 1:**
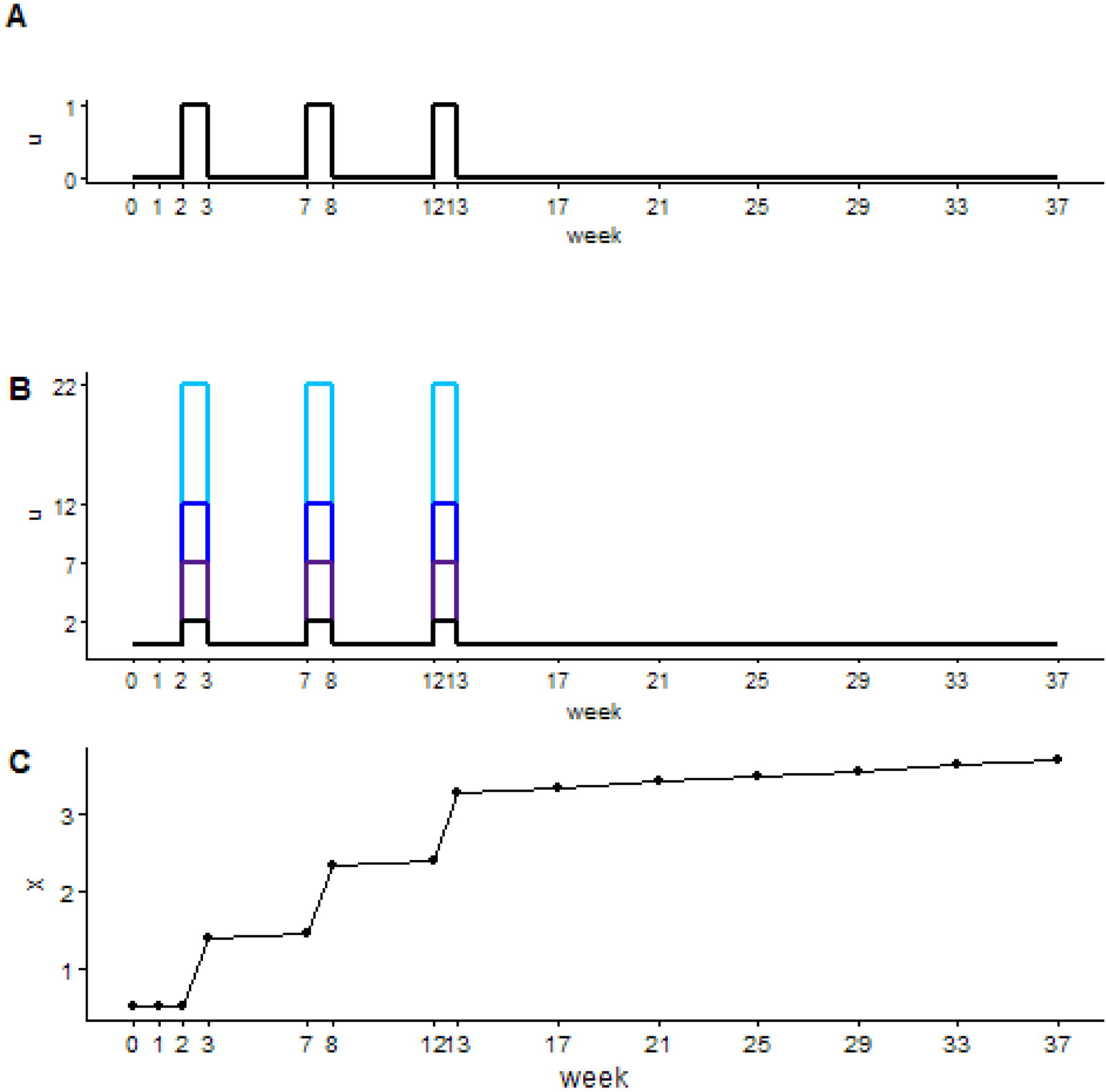
Schematics of the schedules of training and MAL assessments in DOSE. A. Timing of motor training based on the train-wait-train paradigm, in which three 1-week bouts of training are separated by 1-month “wait” periods. B. Weekly doses of training for the four groups. Note that we added 2 hours to the weekly dose in each group to account for the duration of motor assessments– see Methods. C. Example of a simulation showing the changes in the quality of arm movements in the wait-train-wait paradigm modeled with a piece-wise linear model with a positive retention parameter (see Equation 1 in Supplementary Methods). The black dots show the timing of each assessment over the 37 weeks of the DOSE trial.

Using linear models, we previously showed that the changes in MAL due to training in DOSE was dose-dependent^9^. While such “fixed” regression models can predict response to interventions or recovery^9, 10, 12^, they cannot simultaneously account for both the changes in outcomes during training (when an increase in performance is expected) and outside of training for individual patients (where either a decrease in performance due to forgetting or an increase in performance due to self-training is expected). Parsing the repeated measure data to test changes during specific periods (i.e. during training or at different times outside of training) increases variability and decreases power. We need methods that simultaneously use all the data available.

Here, we propose a novel, yet rigorous, statistical modeling framework that accounts for the changes in rehabilitation outcomes during and following training simultaneously. The models are inspired by previous models in motor learning and adaptation^18–26^, in which increases in motor memory are modulated by “learning rate” parameters, and decays in memory are modulated by “retention rate” parameters. Similarly, here, we propose novel piece-wise linear models, in which the increase in MAL during training is modulated by a learning rate parameter and the change in MAL outside of training is modulated by a “retention rate” parameter. In addition, because stroke is characterized by considerable between-subject variability in lesions, recovery, and responsiveness to therapy^27^, subject specific random effects are added to model individual differences in change of performance^28^.

To test for the efficacy, efficiency, and retention of training, the learning and retention parameters are themselves modeled as linear models of experimental variables or participant co-variates, such as the dose of training, the bout of training, time post-training, and the performance post-training. Testing for significance of the slope of these linear parameter models allowed us to test the following hypotheses, derived from motor learning and stroke research, referenced above: 1) The efficacy of training will increase following larger doses of training. 2) The efficiency of training will decrease with larger doses of training. 3) The efficiency of training will decrease for additional bouts of training. 4) Retention will improve with larger doses. 5) Retention will show an initial fast decay followed by a slower decay. 6) The level of performance after training will modulate retention through “self-training”.

## Methods

### The DOSE clinical trial

Detailed inclusion and exclusion criteria of the DOSE trial are described in ^9^. Briefly, participants were included if they were older than 21 years, their score on the impairment UE Fugl-Meyer motor (UEFM) test at baseline was in the 19-60 range out of 66, their cognitive function sufficiently preserved to provide informed consent, and they presented little to no upper extremity sensory impairment or neglect. Each participant signed an informed consent, and the study was approved by the Institutional Review Board of the University of Southern California.

Each of the four doses were distributed over three bouts of training, each separated by 1 month, as shown in Figure 1B. The intervention was based on the Accelerated Skill Acquisition Program (ASAP)^14, 29^. In brief, ASAP includes 1) elements of purposeful and skilled movement execution, including challenging and progressive practice, 2) support for patients’ control or autonomy by choices of specific tasks to be practiced, 3) collaborative problem solving to identify and address movement needs, 4) and encouragement of self-direction in extending practice to community contexts.

Each participant underwent 14 clinical assessments, each including the MAL, the Wolf Motor Function Test (WMFT), and an arm reach performance and choice test, the BART^30^. The assessments were given: 1) twice in the month before training with a two-week interval to assess baseline values; 2) for each of the three 1-week training bouts, in the morning of the first day of training and within 3 days following training; and 3) monthly for 6 months following the last training bout (see Figure 1).

The primary outcome measures in the DOSE trial were the MAL Quality of Movement sub-scale and the WMFT time score. The MAL is a semi-structured interview in which participants are asked to recall and rate the quality of movement of the paretic arm for 28 activities of daily living. It has been used extensively, including in the EXCITE trial, and has good validity and reliability^31^. Here, we analyzed the changes in the MAL Quality of Movement sub-scale, because we previously showed a robust and highly significant dose response for the MAL^9^.

### Modeling the changes in the quality of arm movements during, between, and after training bouts

We built several piece-wise linear models, each to test a specific hypothesis. The learning and retention rates were modeled as linear functions of experimental variables and/or between-subject co-variates (see Table 1; see Supplementary Methods). Specifically, to assess the efficacy of training with increasing dose, the learning rates were functions of dose; the inputs to the model were step inputs of magnitude equal 1, corresponding to the timing of training (Figure 1A; Models 1 in Table 1). To assess the efficiency of training with increasing dose, the learning rates were also a function of dose, but the inputs were modeled as step inputs of magnitudes equal to the weekly dose (Figure 1B; Models 2). To assess the efficiency of training with increasing bouts of training, we modeled the learning rate as a function of increasing weeklong bouts of training (Models 3). To assess the dynamics of retention, we modeled the retention rates as a function of time after training (in two-month increment, Models 4), dose of training (Models 5), and average amount of arm and hand use after training (averaged MAL over 6 months; Models 6).

**Table 1:**
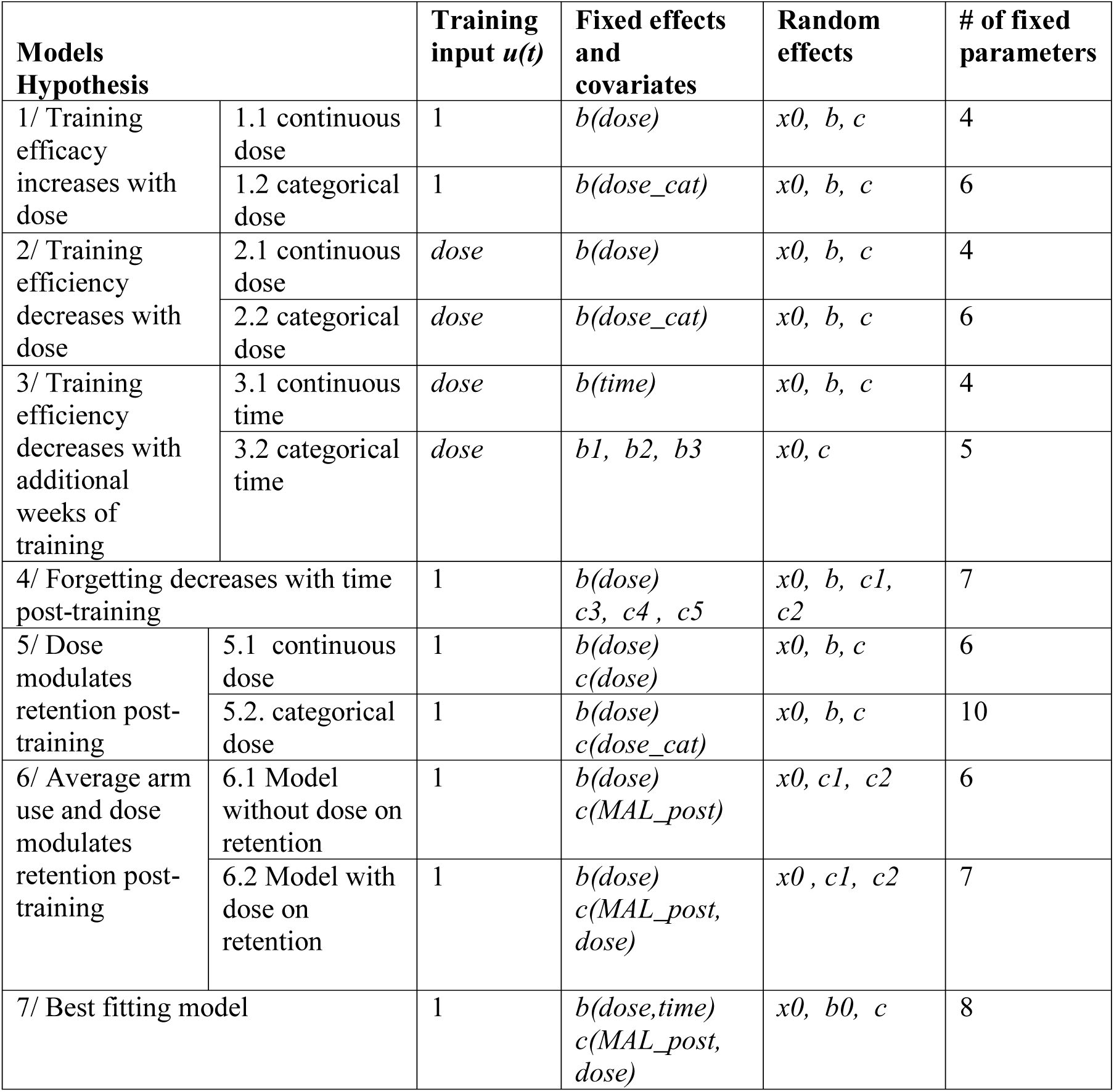
Models selected for all hypotheses tested. Note that the fixed effects X_O_ corresponding to initial MAL are not shown because it was included in all models. See Supplementary Methods for more detailed explanations of each model and Supplementary Tables from 1 to 7 for the values of all fixed effect parameters.

| **Models Hypothesis** |  | **Training input $u(t)$** | **Fixed effects and covariates** | **Random effects** | **# of fixed parameters** |
| --- | --- | --- | --- | --- | --- |
| 1/ Training efficacy increases with dose | 1.1 continuous dose | 1 | $b(dose)$ | $x_0, b, c$ | 4 |
| | 1.2 categorical dose | 1 | $b(dose\_cat)$ | $x_0, b, c$ | 6 |
| 2/ Training efficiency decreases with dose | 2.1 continuous dose | $dose$ | $b(dose)$ | $x_0, b, c$ | 4 |
| | 2.2 categorical dose | $dose$ | $b(dose\_cat)$ | $x_0, b, c$ | 6 |
| 3/ Training efficiency decreases with additional weeks of training | 3.1 continuous time | $dose$ | $b(time)$ | $x_0, b, c$ | 4 |
| | 3.2 categorical time | $dose$ | $b_1, b_2, b_3$ | $x_0, c$ | 5 |
| 4/ Forgetting decreases with time post-training | | 1 | $b(dose)$<br>$c_3, c_4, c_5$ | $x_0, b, c_1, c_2$ | 7 |
| 5/ Dose modulates retention post-training | 5.1 continuous dose | 1 | $b(dose)$<br>$c(dose)$ | $x_0, b, c$ | 6 |
| | 5.2. categorical dose | 1 | $b(dose)$<br>$c(dose\_cat)$ | $x_0, b, c$ | 10 |
| 6/ Average arm use and dose modulates retention post-training | 6.1 Model without dose on retention | 1 | $b(dose)$<br>$c(MAL\_post)$ | $x_0, c_1, c_2$ | 6 |
| | 6.2 Model with dose on retention | 1 | $b(dose)$<br>$c(MAL\_post, dose)$ | $x_0, c_1, c_2$ | 7 |
| 7/ Best fitting model | | 1 | $b(dose, time)$<br>$c(MAL\_post, dose)$ | $x_0, b_0, c$ | 8 |

For most models, we developed both a continuous version, in which the parameters (learning and retention rates) depended linearly on variable of interest (dose, time of training, time post-training, and average MAL post-training, see above), and a corresponding categorical version (see Table 1 and Supplementary Methods). Significance of the fixed slopes for the learning or retention linear models was used to test our hypotheses. Comparison of the fixed effect learning or retention parameters of the categorical models to the linear models helped visualize possible deviation from linearity.

Note that we added 2 hours to the weekly dose in each group to account for the approximate duration of the two sets of assessments (which include WMFT and BART arm reaching and arm choice tests) given immediately before and after each training week. The weekly doses of training were therefore equal to 2, 7, 12, and 22 hours. To account for the large variability in initial arm and hand use, we modeled MAL as a function of initial *MALinit* at baseline (median of MAL at 0, 1, and 2 weeks) in all models. Incorporating *MALinit* as a covariate was significant for all models (p < 0.01).

## Results

### Qualitative predictions from the best fitting model

Figure 2A shows both actual MAL data and the fits of the best fitting model (Model 7 in Table 1) for all 41 participants over the 37 weeks of the DOSE trial.. The model well accounts for changes in MAL during both training and following training for 40 subjects (one subject exhibited poor fit due to highly variable measured MAL; see third row second column). Figure 2B shows the model fits re-arranged by nominal doses. Visual inspections of the plots allow the following general observations: 1) Larger initial increase in MAL for larger doses (increased efficacy of training); 2) Diminishing returns for larger doses (decreased dose efficiency); 3) Diminishing returns for more training weeks (decreased time efficiency); 4) Decay of MAL post-training for larger doses; 5) Further increase in MAL in the months following training for participants with high MAL post-training; and 6) in 0 dose group, increase in MAL during and following training. We confirm these observations in the analyses below using the models of Table 1.

**Figure 2:**
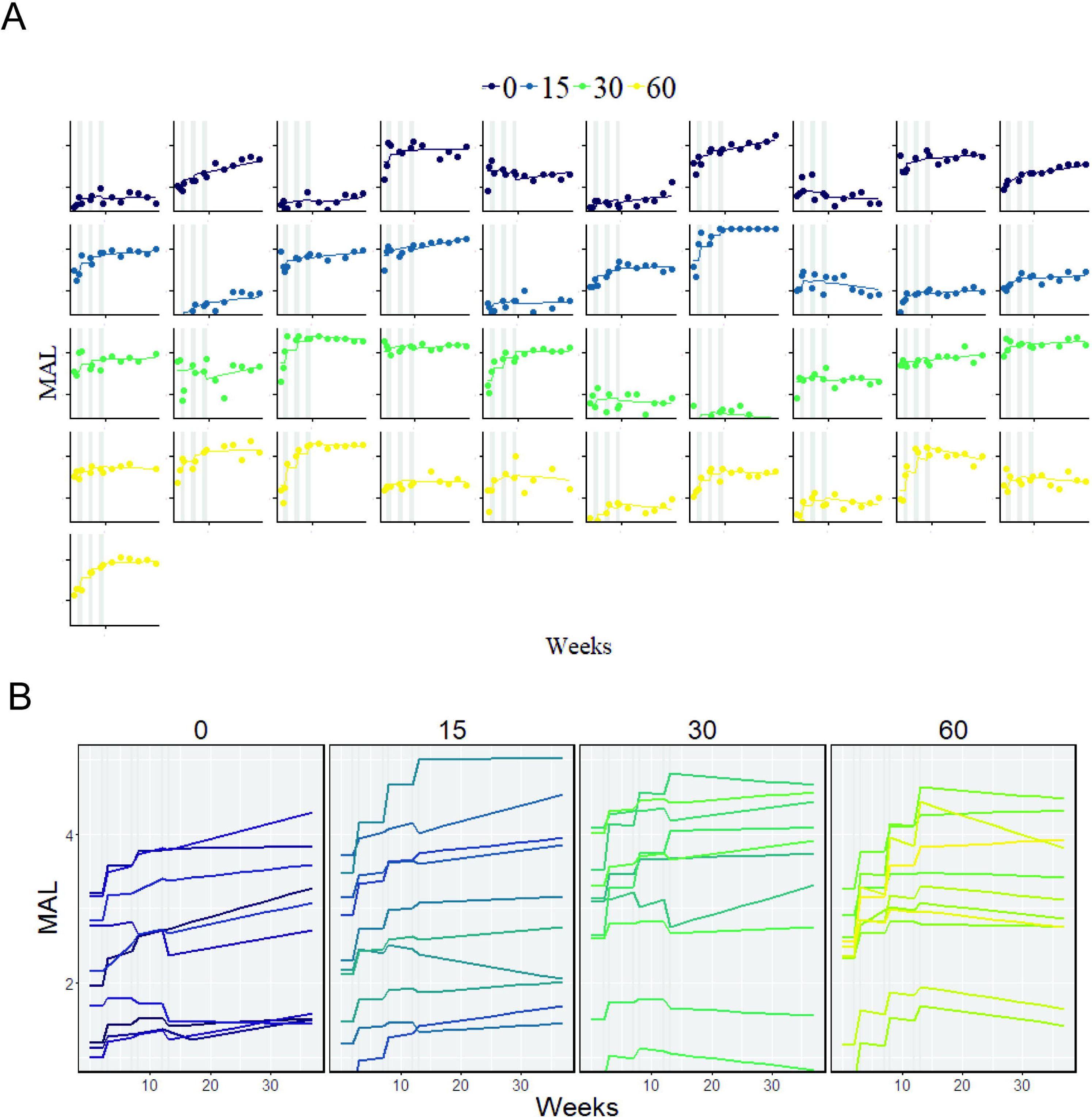
Quality of movement as measured by the MAL over the 37 weeks of the DOSE trial, assessed before, during, between, and following the three training bouts, and an example of model fit to the data, using the best fitting model (Model 7 in Table 1). A. Data (dots) and model fit (lines) for all 41 subjects. Gray vertical lines show the three week-long bouts of training. Note the overall excellent fit of the model to all subjects (except for 2^nd^ subject in 30 hour dose, who exhibited large MAL variability). B. Individual models re-arranged by nominal doses. Note the following: 1) Larger increase in MAL during training for larger doses (increased dose efficacy); 2) Diminishing returns for larger doses (decreased dose efficiency); 3) Diminishing returns for more training weeks (decreased training bout efficiency); 4) Decrease in MAL post-training for larger doses; 5) Further increase in MAL post-training for high average MAL post-training; and 6) in 0 dose group, increase in MAL. These observations were all confirmed statistically see text.

### Increasing the dose increases the efficacy of training

Increasing the dose of training increases the efficacy of training, as shown by the positive slope between dose and the learning rate in Model 1.1 (Table 1; Figure 3A: slope parameter *b.dose* = 0.008; p = 0.032; see Supplementary Table 1 for values of additional model parameters). Inspection of the coefficients of the categorical dose Model 1.2 shows that this increase in efficacy is mostly driven by the 60 hour dose (*b.dose_hour 60* greater than *b.Intercept* p = 0.011; see Supplementary Table 1). Because the initial MAL is larger in the 30 hour dose groups (ANOVA, p = 0.01, 0 dose 2.68 ± 0.57:; 15 hour dose 2.89 ± 0.64, 30 hour dose 3.67 ± 0.94 and 60 hour doses 2.84 ± 0.57), participants in this group appeared to have benefited less from training. In addition, note how the 0 hour group shows an increase in the MAL.

**Figure 3:**
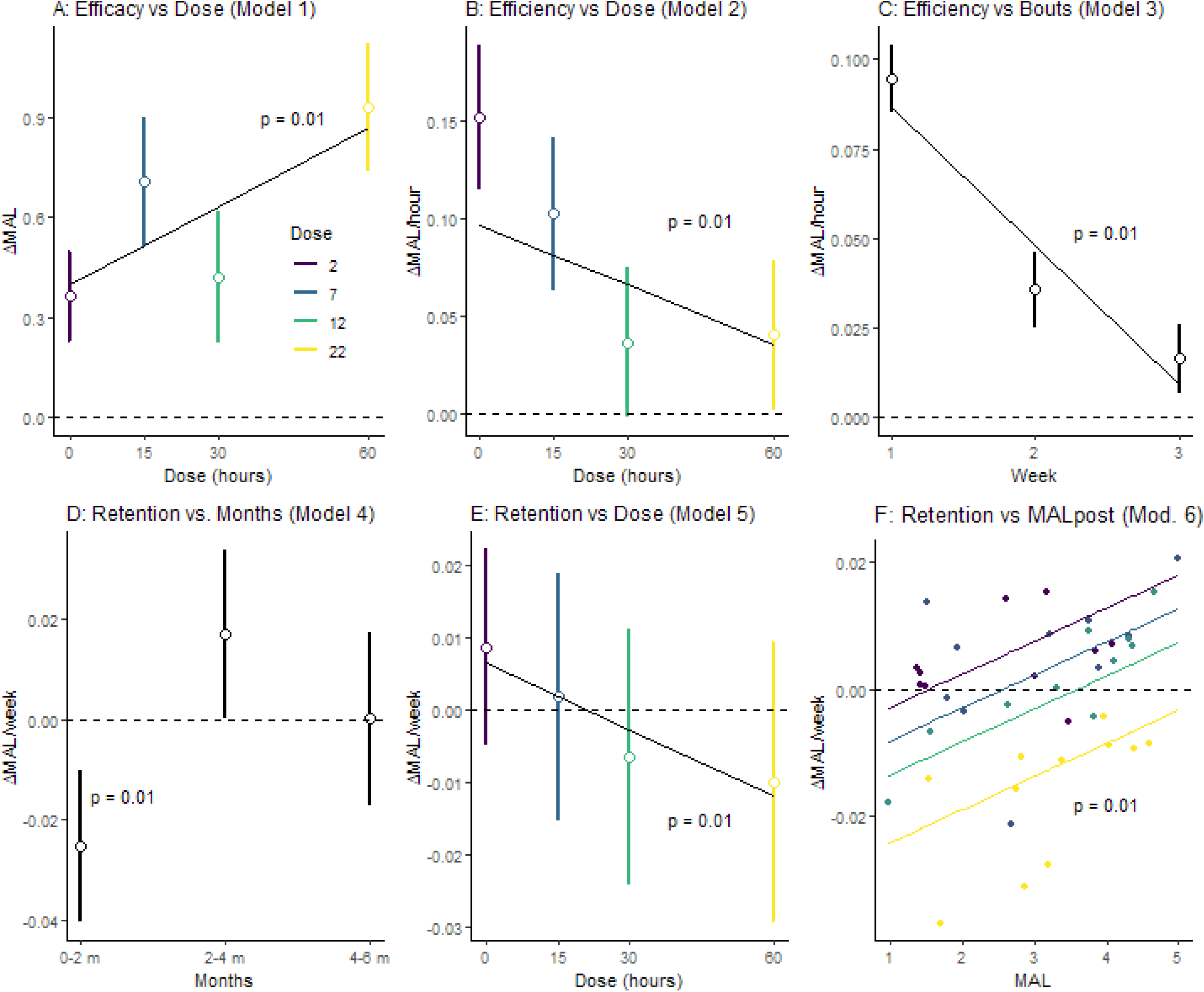
Fixed effect parameters used to test all six hypotheses. A. Effect of dose on the MAL during training (dose efficacy; Models 1 in Table 1). B. Effect of increasing the dose on gain in MAL per hour of training (dose efficiency; Models 2). C. Effect of increasing the number of weeks of training on gain in MAL per hour (duration efficiency; Models 3). D. Effect of time on retention: Retention rate in the 6 months following training as a function of the number of months post-training (0-1 months, 2-3 months, and 3 to 5 months) (Model 4). E. Effect of dose on retention (Models 5). F. Effect of average MAL post training on gain in MAL post training (Models 6). The thick dark line shows the retention as a function of average MAL post training in Model 6.1. The dashed vertical line indicates the threshold (= 3.4) in average MAL post above which the MAL continues to increase post-training. The colored line shows the retention as a function of average MAL post for the different doses and the colored dots show the individual retention rates (mixed effects) in Model 6.2.

### Increasing the dose decreases the efficiency of training

We then studied the efficiency of training, that is, the gain in MAL per hour for increasing doses (Models 2 in Table 1). As predicted, increasing the dosage of training decreases the efficiency of training, as shown by the negative slope (Model 2.1; Figure 3B; parameter *b.dose* = - 0.001; p = 0.012). Inspection of the coefficients of the categorical model (See Supplementary Table 2) shows that efficiency largely decreases from 0 hour until 30 hours and is then similar for 30 and 60 hours (0 hour group, efficiency per week of training: 0.050, p = 0.002; decrease in efficiency compared to the 0 hour: 30 hours: −0.038, p = 0.019; 60 hour, −0.037 p = 0.022. (Note because we considered that the 0 hour group received 2 hours of motor training, efficiency could be computed for the 0 hour group as well – see above).

### Increased training duration decreases the efficiency of training

As was the case for increasing doses, the efficiency of training decreases with the number of additional weeks of training (Model 3.1; Figure 3C; *bt.weeks*= −0.013; p < 0.0001, see Supplementary Table 3). The categorical model (Model 3.2) shows a more than two-fold reduction in gain for the second week and almost five-fold reduction for the third week compared to the first week (see Supplementary Table 3). Interestingly, the *b3* coefficient (learning rate for the third week) in the categorical model 3.2 is not significantly different from zero (p = 0.15), showing that across doses, the third bout of training has very little effect on changes in MAL during training (note however that because we did not vary the number of weeks in the DOSE trial, we cannot determine how the third bout influences retention).

### Decay following training is fast initially but slows down within 2 months post-training

In motor learning research, it is commonly observed that forgetting following training is initially fast and then the rate of forgetting gradually declines. We thus developed a model in which retention rates (in units of change of MAL per week) are estimated over three intervals of 2 months each during the six months after training (Model 4 in Table 1). Results show that the retention parameter *c1* in the 2 months following training is largely negative, but retention is not different from zero in the next two 2-month periods although there is a non-significant trend upward for the next two months (Figure 3D; *c1* = −0.025, p = 0.002; *c2* = 0.017, p = 0.50; *c3* = 0.0001, p = 1; see Supplementary Table 4).

### Retention following training is negatively modulated by the dose of training

Following training, we found an unexpected increase in forgetting for larger doses, as seen by the negative slope between dose and the change in MAL between sessions (Figure 3E; Model 5 in Table 1, parameter *c.dose_cont* = −0.001; p = 0.01; see Supplementary Table 5). The model with categorical doses (Model 5.2 in Table 1) shows that the effect of dose on retention is negative and significantly different from 0 for 30 and 60 hour doses (*c.dose_hour30* = −0.015; p = 0.023, *c.dose_hour60* = −0.019; p = 0.012), but not for 15 hours (p = 0.28). Because of the decrease in the MAL following training in the large dose groups, the dose response curve observed during training (Figure 3A) does not hold from pre-training to 6 months post-training. Indeed, there is no difference in response across doses in the differences in the initial MAL before training with the last MAL after training (medians of the first three MAL in assessments 1, 2, and 3, and medians of the last three MAL in assessments 12, 13, and 14; ANOVA, p = 0.27).

### A threshold determines increase versus decrease in the quality of arm movement following training

As seen above, the higher the dose of training, the more the forgetting overall. However, as we have previously proposed^13, 17^, and as appears to be indicated through visual observation in Figure 2, higher levels of spontaneous use following training may reduce decay, or even further increase use. Here, we therefore test the hypothesis that retention in the 6 months following training depends on the average MAL *MALpost* during these 6 months, with greater MAL leading to increased retention. Figure 3F shows that this is indeed the case: retention increases with the average MAL in the six months post training (Model 6.1 slope = 0.0047, p = 0.02; see Supplementary Table 6). Because of the negative intercept in the linear model (−s0.016, p = 0.019), there is a threshold for spontaneous increase in MAL post training. Thus, on average, across doses, if the *MALpost* is above a threshold of 3.4 (determined by 0.0047 * *MALpost* −0.016 = 0; see vertical dashed line, Figure 3F), the retention parameter is positive, that is, MAL keeps increasing following additional training (besides the additional motor assessments in the 6 month follow-up period).

A model that includes both *MALpost* and dose on the retention parameter (Model 6.2) shows that for a smaller dose (0, 15, and 30 hours), the retention parameter is negative for smaller values of MAL post and positive for larger value (see Figure 3F and Supplementary Table 6). However, because increasing dose leads to lower retention (see above), the threshold above which the MAL increases following training is dose-dependent, with the counter-intuitive results that a smaller average *MALpost* can be sufficient to increase MAL following training for small doses but not for the largest dose (see Figure 3F). For instance, for the 15 hour dose, the threshold is approximately 2.5. In contrast, for the 30 hour it is just under 4. For the 60 hour dose, retention is negative for all values of average MAL post. Figure 2B shows such a negative retention parameter for nine participants out of 11 in the 60-hour group.

## Discussion

Increasing the efficacy of training (“total gain due to training”), the efficiency of training (“gain per unit time of training”), and the long-term retention (“durability of gains”) following motor training after stroke form the cornerstones of neurorehabilitation. Twenty years ago, Fuhrer and Keith proposed that “the effectiveness and efficiency of learning-oriented practices will likely be enhanced by well-formulated investigations grounded in available learning theory and research”^32^. Here, using a combination of novel piece-wise linear models with mixed effects and a novel rehabilitation practice design, we were able to dissociate the effects of dose and duration of training on the efficacy, efficiency, and retention of neurorehabilitation on arm and hand use.

Our findings show that large doses of practice increase the MAL from pre-to post-training (Figure 3A). This result reproduces the previously reported dose-response relationship for the MAL in the DOSE trial, which was only determined from the changes due to the three bouts of training^9^. Here, fitting mixed-effects piece-wise models that include all the data, in particular the 1-month wait periods between the training bouts, the learning rates of the model represent the gain in MAL per week during training. Thus, the dose versus learning rates plot also represents a dose-response curve (Figure 3A).

In contrast, a model with inputs that scale to the dose of training showed a clear decrease in efficiency with additional hours of training (Figure 3B). Each hour of training in the 60-hour dose group is about two times less effective in increasing the MAL than an hour in the 15-hour dose. Similarly, there is a strong decrease in the efficiency of weeks of training: the first week has greater efficiency than the second, itself greater than the third week (Figure 3C). The third week is more than five times less efficient than the first. Such decreases in efficiency are consistent with the well-known power law of motor learning, where each additional unit of practice yields a smaller gain.

Across doses, retention following training resembles exponential-like decay (Figure 3D), because it is initially fast in the two first month following training before tapering back near baseline in the next 4 months. Such decay-like forgetting is seen in motor adaptation, which is defined as the ability to gradually modify motor commands in order to compensate for changes in our body and in the environment^33^.

Contrary to our hypothesized effect of increasing dose on retention, we found that large doses increase decay in the six months post-training (Figure 3E). Because of this dose-dependent decay, there was no dose-response between baseline and end of follow-up. A plausible explanation for this apparent negative effect of dose is poor transfer. While ASAP emphasized self-management skills and the development of self-efficacy for meaningful arm use, it did not include a transfer package^34^, which consists of additional time at the end of every training session with review of daily arm use and problem solving to overcome perceived barriers to arm use at home. Inclusion of such transfer package in CIMT (constraint-induced movement training) lead to large behavioral changes in real-world arm function. ^34^

However, our results suggest that spontaneous arm use post-training modulated retention. We previously postulated the existence of a threshold in which arm and hand use in daily activities acts as “self-training” and reinforces performance, which then further reinforces use in a virtuous cycle^13, 15, 17^. In line with this previous study, we showed that retention is positively modulated by the average MAL post-training (Figure 3F). Across doses, the threshold for MAL is ∼3.4. Thus, if MAL post-training is relatively high, then the MAL will keep increasing following training. Such an effect is more pronounced in the low dose groups, as can be seen in Figure 2B for a number of participants, because in these groups, the negative effects of dose on retention is less strong.

Finally, as noted the 0 hour group shows significant increase in the MAL during and post-training. In addition to the MAL, each assessment consisted of arm reaching tests comprised with approximately 200 movements with the more affected arm^11^, in addition to the WMFT test (in which subjects perform goal-directed arm and hand movements). Thus, theassessments may have resulted in motor training. It is therefore possible that self-training was complemented by a certain amount of “forced” training during the 6 assessments in the follow-up phase.

### Limitations

There are two primary limitations with our study. First, because of the relatively small number of participants, the groups are not well balanced for initial level of MAL. In particular, because the initial MAL is larger in the 30 hour dose group, participants in this group appeared to have benefited less from training. This affected the dose-dependent results for this group (see Figure 3A). For instance re-running the linear dose efficacy Model 1.1 without the 4 participants with *MALinit* > 3.5 (3 participants in the 30 hour group and 1 participant in the 15 hour group) improved the significance of the slope from p = 0.032 to p = 0.019. Thus, greater number of participants are needed. However, given the extreme difficulty of collecting a large number of repeated measures with clinical tools in the laboratory (in this study, participants were required to make a total of 26 visits each), wearable sensors that can measure arm and hand use, such as accelerometers^35^ or the manumeter^36^, should be considered in future work.

Second, and related, it may be argued that our number of participants was too small to test six hypotheses with sufficient power. However, we believe that our results are valid because: 1) our analyses were pre-planned based on hypotheses derived from previous research, 2) all our results, except for the unexpected dose-dependent decay, matched our hypotheses, 3) we reproduced two previous results obtained with different methods (the dose response relationship^9^ and the post-training arm use threshold^17^, 4) the six related continuous models contained a maximum of 7 fixed effect parameters estimated from a total of 559 data points for 41 participants (15 data points were missing overall), and finally 5) the power of mixed effect models increases both with number of subjects (41 here) and number of repeated measures (14 here) for each subject^37^.

## Conclusions: implications for clinical practice

Our results paint a contrasted view of the effect of motor therapy on arm use in the chronic stage post-stroke: although training increases arm use, these gains due to training are often lost following training, notably for larger doses. Thus, large doses may not be highly effective overall, in part because the efficiency of each training hour quickly decreases. However, the further increase of the MAL post-training for participants with high average MAL post-training in the lower dose groups casts hope for developing effective and efficient personalized rehabilitation methods.

Specifically, our results suggest three methods to maximize the efficacy and efficiency of training in the chronic stage post-stroke: First, motor training in the chronic stage should be given in relatively small bouts distributed over months. Such strategy would not only maximally increase arm and hand use for each hour of training (because smaller doses of training are more efficient than larger doses), but also maintain the gains (because smaller doses lead to less decay). This training strategy is consistent with one of the most robust effects of learning theory, the so-called “spacing effect”, according to which the spacing of presentations strongly increases retention compared to massed presentations^24, 38–40^. Second, these small bouts of training should contain intensive movement training. The number of movements in the 0 dose group of the present study is approximately 200 movements. We previously showed that most gains in reaching movements were achieved with approximately 300 movements^41^. This can be contrasted to the amount of training given in a single therapy session in clinical practice (32 movements on average per session^42^), which is probably insufficient to have a large effect on arm and hand use. Delivering such large number of training movements will probably be well suited via technological devices that train the movements, such as the BART system^30^.

Third, our results suggest a personalized dosage duration of therapy based on arm and hand use measurements: We showed that, on average, for individuals with MAL above threshold of approximately 3.4 post-training, motor therapy could be stopped, as self-training in daily activities will continue to increase arm use. Thus, perhaps all these above-threshold individuals need is a self-guided home program or a customized transfer package. For individuals with MAL below this threshold, decay in arm use follows therapy, on average. Thus, for these individuals, our results suggest that there is a need for a program to build confidence in arm use and develop strategies for overcoming barriers to arm use at home, so that they can more effectively engage in self-training.

## Acknowledgements

We thank Bokkyu Kim and Sujin Kim for their help in pre-processing the data.

## Supplementary Methods

### Modeling the dynamics of recovery during and after rehabilitation

#### Mixed effects dynamical model to model the change in MAL during and outside therapy

Here, we developed dynamical models to account for the time course of the changes in the MAL in response to discrete bouts of training with different doses. In addition, mixed effects models were added to account for the high between-individual variability in lesion, impairment, and responsiveness to therapy. In a previous reach training study with participants with chronic stroke (Park and Schweighofer, 2017), we used non-linear mixed effects models to dissociate improvement in performance due to learning from highly variable decrease in performance presumably due to fatigue. Dynamical models with mixed effects have been used widely to model the action of medicines as part of the clinical trial drug development process (Sheiner and Beal, 1981).

Our models are based on simple first order linear equations with a time step of 1 week, which corresponds to the duration of each bout of training. Let *x*(*t*) be the MAL at time t, with t in increments of days, with input *u*(*t*). We developed two main classes of models. In models aimed at testing the efficacy of training, *u*(*t*) represents the timing of training (Figure 1A). In models aimed at testing the efficiency as a function of dose for instance, *u*(*t*) represents both the dose and timing of training (Figure 1B). In both cases, *u*(*t*) = 0, before, between, and after training bouts. Note that the nominal doses were 0, 15, 30, and 60 hours per group, distributed over the three training weeks. However, in the analyses, we added 2 hours per training weeks to account for the approximate duration of motor tests (WMFT and BART) before and after training (hence 2, 7, 12, and 22 hours per week). To isolate the effect of the training input *u*(*t*), we further assumed that forgetting was smaller than learning during training. Thus, the simplest first order dynamical model of the changes in MAL for a single subject is given by:

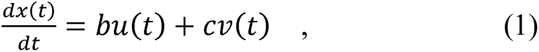

where 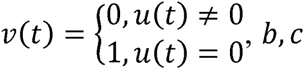 represent constant “learning” and “retention” rates, respectively.

We did not constrain *b* or *c* to be positive or negative, although we expected *b* to be positive as we previously showed a positive relationship between dose and changes in MAL (Winstein et al., 2019).

Given the definition of *u*(*t*) and *v(t)*, the solution to Eqn. (1) is a piece-wise linear equation. Thus, although the response during each bout and following each bout is linear, the overall solution *x* = *f*(*x_0_, b, c, t*) is non-linear, where *x*_0_ is the median MAL in the three test before training. We then updated this nonlinear function to include mixed effects. In its general form, the model is given by:

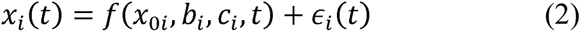

where *x_0i_, b_i_, c_i_* are the mixed effect parameters for individual *i*, *ϵ*_i_(t) is the residual, assumed to be normally distributed with constant variance (Lindstrom, 1990).

Model selection in general, and inclusion of random effects in particular, was based on the log-likelihood ratio test for nested models and, for non-nested models on Akaike and Bayesian Information Criteria (AIC and BIC), which provide measures of the quality of fit by minimizing fitting error and penalizing the number of model parameters (See Supplementary Methods for details and rationale). A list of models tested is given in Table 1.

#### Modeling efficacy and dose-efficiency of training

##### Continuous dose models

To estimate the dose-efficacy of training (the overall effect of training), we let *u*(*t*) = 1 during the intervention periods as shown in Figure 1A. To estimate the dose-efficiency of training (the effect of training per hour), we let *u*(*t*) to be the number of treatment hours corresponding to the dose, as shown in Figure 1B. To investigate how training efficacy and efficiency depends on dosage of training, the mixed effects of *b_i_* are given by:

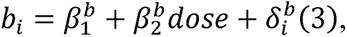

where *dose* is a continuous variable, 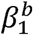 and 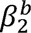 are the fixed effects relating dose to the mean of *b* in the population; is the value of the random deviation of the ith subject from the population mean value. It is assumed that 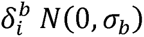 where *σ_b_* reflects the inter-subject variability in *b*.

Similarly, to investigate how retention depends on dosage of training, the mixed effects of *c_i_* are given by:

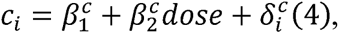

##### Categorical dose models

To study how efficacy and efficiency depends on the four doses, we also developed models in which dose of training is taken as categorical variable. The formula becomes

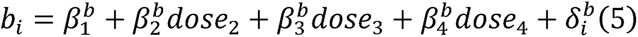

where *dose_i_* is 1 for dose *j*, 0 otherwise. The formula for *c_i_* follows the same structure. In this, and all other models, the initial MAL is included as

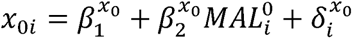

Where it is assumed that 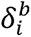, where 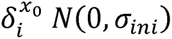,, where *σ_ini_* reflects the inter-subject variability in initial 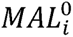, the initial MAL score of participant *i*.

#### Modeling time-efficiency

To study the effect of training in successive bouts, we assumed b can take different values during training. In the models with continuous effect of time, we considered a linear dependence of *b* and *c* on training period *k*(*k* = 1, 2,3), namely

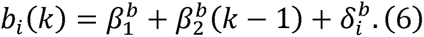

A similar model for categorical effect of training was developed.

#### Modeling decay post-training

To study the effects of training on retention, we develop two additional models. In the first model, we tested whether the MAL returned to its initial baseline or was maintained following training. For this, we sliced the 6 months post-training in three equal slices of 2 months, and defined three post-training decay parameters for each 2 months period. For each, we assume a dose relationship such that 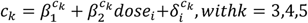:

In the second model, we test the hypothesis that decay in the six months post-training is due to both dose and the average MAL in the six months post-training. For this, we model

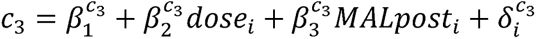

#### Model fitting

Model fits were performed using the *nlme()* function in R, which uses maximum likelihood to estimation (Pinheiro et al., 2017). For categorical models, the 0 dose group was used as the reference group, such that the group effects for the other three dose groups are represented with respect to the 0 group. For these models, the R function *intervals()* was used to construct the standard errors for the fixed parameters (in Supplementary Tables).

Except for the best fitting model, the fixed effects of each models were selected to test specific hypotheses. In addition, because the parameters are estimated from 559 data points (the ideal number is 41 * 14 = 574 points; 15 data points were missing), the models appear sufficiently small not to over-fit the data. For instance, the best fitting model (Model 7) contains 8 fixed effect parameters and 3 random effect parameters (random parameter variance). In typical model development, over-fitting is typically best avoided with cross-validation; this cannot be performed with random effects, however, because cross-validation requires “out-of-sample” predictions, which cannot be easily calculated when random effects are present. However, it has been shown that AIC applied to mixed models is equivalent to leave-one-cluster-out cross-validation. Thus model selection via both AIC and BIC supports the proposed models. Additional statistical analyses were performed in *R* 3.4.0. Statistical significance was predetermined at p < 0.05.

Note on doses included in the model: the nominal doses are 0, 15, 30, and 60 hours of ASAP therapy. Because pre- and post-training tests lasted for approximately 1 hours, two hours per training bouts were added to the actual dose of training in the model. Thus, whereas the actual doses of therapy were 0, 5, 10, and 20 hours per week, the doses used in the model were 2, 7, 12, and 22 hours (See Figure 1B). Note that the addition of a non-zero dose for the control group allows defining a learning rate parameter for this group, and therefore allows us to compare learning rate in other groups to this reference group.

---

**Supplementary Table 1:** Dose efficacy Model 1.1 for continuous dose and 1.2 for categorical doses.

|  | Cont. Dose | Cat. Doses |
| --- | --- | --- |
| x0.(Intercept) | 0.184<sup>*</sup><br>(0.093)<br>p = 0.048 | 0.191<sup>*</sup><br>(0.094)<br>p = 0.042 |
| x0.iniMAL | 0.954<sup>***</sup><br>(0.036)<br>p = 0.000 | 0.951<sup>***</sup><br>(0.036)<br>p = 0.000 |
| b.(Intercept) | 0.118<sup>*</sup><br>(0.048)<br>p = 0.016 | 0.121<sup>*</sup><br>(0.053)<br>p = 0.024 |
| b.dose_cont | 0.008<sup>*</sup><br>(0.004)<br>p = 0.032 |  |
| b.dose_cat15 |  | 0.114<br>(0.075)<br>p = 0.127 |
| b.dose_cat30 |  | 0.019<br>(0.075)<br>p = 0.803 |
| b.dose_cat60 |  | 0.188<sup>*</sup><br>(0.073)<br>p = 0.011 |
| Observations | 557 | 557 |
| Log Likelihood | -261.199 | -259.527 |
| Akaike Inf. Crit. | 544.398 | 545.054 |
| Bayesian Inf. Crit. | 591.946 | 601.247 |
| *Note:* | <sup>*</sup>p<0.05; <sup>**</sup>p<0.01; <sup>***</sup>p<0.001 |  |

**Supplementary Table 2:** Dose efficiency Models 2.1 for continuous dose and Model 2.2 for categorical doses.

|  | Cont. Dose | Cat. Doses |
| --- | --- | --- |
| x0.(Intercept) | 0.166<br>(0.096)<br>p = 0.083 | 0.168<br>(0.098)<br>p = 0.087 |
| x0.iniMAL | 0.968***<br>(0.037)<br>p = 0.000 | 0.962***<br>(0.038)<br>p = 0.000 |
| b.(Intercept) | 0.034***<br>(0.007)<br>p = 0.00000 | 0.050**<br>(0.015)<br>p = 0.002 |
| b.dose_cont | -0.001*<br>(0.0004)<br>p = 0.012 |  |
| b.dose_cat15 |  | -0.016<br>(0.016)<br>p = 0.322 |
| b.dose_cat30 |  | -0.038*<br>(0.016)<br>p = 0.019 |
| b.dose_cat60 |  | -0.037*<br>(0.016)<br>p = 0.022 |
| Observations | 557 | 557 |
| Log Likelihood | -267.756 | -263.789 |
| Akaike Inf. Crit. | 557.512 | 553.578 |
| Bayesian Inf. Crit. | 605.060 | 609.772 |
| Note: | *p<0.05; **p<0.01; ***p<0.001 |  |

**Supplementary Table 3:** Efficiency of training weeks Model 3.1 for continuous week of training and Model 3.2 for categorical weeks of training.

|  | Cont. weeks | Cat. weeks |
| --- | --- | --- |
| x0.(Intercept) | 0.168<br>(0.126)<br>p = 0.183 | 0.165<br>(0.126)<br>p = 0.191 |
| x0.iniMAL | 0.935***<br>(0.049)<br>p = 0.000 | 0.935***<br>(0.049)<br>p = 0.000 |
| b0 | 0.029***<br>(0.003)<br>p = 0.000 |  |
| bt | -0.013***<br>(0.003)<br>p = 0.00000 |  |
| b1 |  | 0.031***<br>(0.004)<br>p = 0.000 |
| b2 |  | 0.012**<br>(0.004)<br>p = 0.005 |
| b3 |  | 0.006<br>(0.004)<br>p = 0.153 |
| Observations | 557 | 557 |
| Log Likelihood | -288.013 | -287.491 |
| Akaike Inf. Crit. | 592.027 | 592.981 |
| Bayesian Inf. Crit. | 626.607 | 631.884 |
| Note: | * p<0.05; ** p<0.01; *** p<0.001 |  |

**Supplementary Table 4.** Retention with time, Model 4

|  |  |
| --- | --- |
| x0.(Intercept) | 0.142 |
|  | (0.093) |
|  | p = 0.127 |
| x0.iniMAL | 0.958<sup>***</sup> |
|  | (0.036) |
|  | p = 0.000 |
| b.(Intercept) | 0.157<sup>**</sup> |
|  | (0.049) |
|  | p = 0.002 |
| b.dose_cont | 0.005 |
|  | (0.003) |
|  | p = 0.148 |
| c1 | -0.025<sup>**</sup> |
|  | (0.008) |
|  | p = 0.002 |
| c2 | 0.017<sup>*</sup> |
|  | (0.009) |
|  | p = 0.050 |
| c3 | 0.0001 |
|  | (0.009) |
|  | p = 0.995 |
| Observations | 557 |
| Log Likelihood | -254.921 |
| Akaike Inf. Crit. | 545.842 |
| Bayesian Inf. Crit. | 623.649 |
| Note: | * p<0.05; ** p<0.01; *** p<0.001 |

**Supplementary Table 5.** Effect of dose on Retention Model 5.1 for continuous dose and Model 5.2 for categorical doses

|  | Cont. Dose | Cat. Doses |
| --- | --- | --- |
| x0.(Intercept) | 0.187<sup>*</sup><br>(0.094)<br>p = 0.048 | 0.185<sup>*</sup><br>(0.094)<br>p = 0.050 |
| x0.iniMAL | 0.954<sup>***</sup><br>(0.037)<br>p = 0.000 | 0.955<sup>***</sup><br>(0.037)<br>p = 0.000 |
| b.(Intercept) | 0.065<br>(0.058)<br>p = 0.258 | 0.065<br>(0.058)<br>p = 0.260 |
| b.dose_cont | 0.014<sup>**</sup><br>(0.004)<br>p = 0.002 | 0.014<sup>**</sup><br>(0.004)<br>p = 0.002 |
| c.(Intercept) | 0.008<br>(0.005)<br>p = 0.076 | 0.009<br>(0.005)<br>p = 0.088 |
| c.dose_cont | -0.001<sup>**</sup><br>(0.0004)<br>p = 0.010 |  |
| c.dose_cat15 |  | -0.007<br>(0.006)<br>p = 0.281 |
| c.dose_cat30 |  | -0.015<sup>*</sup><br>(0.007)<br>p = 0.023 |
| c.dose_cat60 |  | -0.019<sup>*</sup><br>(0.007)<br>p = 0.012 |
| Observations | 557 | 557 |
| Log Likelihood | -257.782 | -257.209 |
| Akaike Inf. Crit. | 541.564 | 544.418 |
| Bayesian Inf. Crit. | 597.758 | 609.256 |
| Note: | <sup>*</sup>p<0.05; <sup>**</sup>p<0.01; <sup>***</sup>p<0.001 |  |

**Supplementary Table 6.** Effect on retention of average MAL post-training, Model 6.1, and Model 6.2 of average MAL post-training and continuous doses

|  | MALpost | MALpost + Dose |
| --- | --- | --- |
|  | (1) | (2) |
| x0.(Intercept) | 0.208<sup>*</sup> | 0.209<sup>*</sup> |
|  | (0.093) | (0.094) |
|  | p = 0.025 | p = 0.027 |
| x0.iniMAL | 0.945<sup>***</sup> | 0.945<sup>***</sup> |
|  | (0.036) | (0.037) |
|  | p = 0.000 | p = 0.000 |
| b.(Intercept) | 0.150<sup>**</sup> | 0.066 |
|  | (0.050) | (0.058) |
|  | p = 0.004 | p = 0.254 |
| b.dose_cont | 0.006 | 0.014<sup>**</sup> |
|  | (0.003) | (0.004) |
|  | p = 0.081 | p = 0.002 |
| c.(Intercept) | -0.016<sup>*</sup> | -0.006 |
|  | (0.007) | (0.007) |
|  | p = 0.019 | p = 0.417 |
| c.dose_cont |  | -0.001<sup>**</sup> |
|  |  | (0.0004) |
|  |  | p = 0.004 |
| c.MALpost | 0.0047<sup>*</sup> | 0.005<sup>**</sup> |
|  | (0.002) | (0.002) |
|  | p = 0.020 | p = 0.008 |
| Observations | 557 | 557 |
| Log Likelihood | -259.009 | -254.975 |
| Akaike Inf. Crit. | 544.017 | 537.951 |
| Bayesian Inf. Crit. | 600.211 | 598.467 |
| Note: | <sup>*</sup> p<0.05; <sup>**</sup> p<0.01; <sup>***</sup> p<0.001 |  |

**Supplementary Table 7.** Best Model 7, as indicted by the lowest BIC, found via combination of the continuous covariates in the other models.

|  |  |
| --- | --- |
| x0.(Intercept) | 0.148 |
|  | (0.093) |
|  | p = 0.112 |
| x0.iniMAL | 0.944*** |
|  | (0.036) |
|  | p = 0.000 |
| b0.(Intercept) | 0.220*** |
|  | (0.065) |
|  | p = 0.001 |
| b0.dose_cont | 0.013** |
|  | (0.004) |
|  | p = 0.002 |
| bt | -0.175*** |
|  | (0.034) |
|  | p = 0.00000 |
| c.(Intercept) | -0.001 |
|  | (0.007) |
|  | p = 0.912 |
| c.dose_cont | -0.001** |
|  | (0.0004) |
|  | p = 0.004 |
| c.MALpost | 0.005** |
|  | (0.002) |
|  | p = 0.008 |
| Observations | 557 |
| Log Likelihood | -242.180 |
| Akaike Inf. Crit. | 514.361 |
| Bayesian Inf. Crit. | 579.199 |
| Note: | * p<0.05; ** p<0.01; *** p<0.001 |

